# Mesoglea biogenesis reveals a cryptic aboral valve for pressure regulation in cnidarian morphogenesis

**DOI:** 10.1101/2025.05.26.656114

**Authors:** Soham Basu, Petrus Steenbergen, Florian Gabler, Alexandre Paix, Paolo Ronchi, Gleb Bourenkov, Thomas Schneider, Jonas Hellgoth, Anna Kreshuk, Suat Özbek, Aissam Ikmi

## Abstract

Cnidarians are classically defined by a single oral opening, a hallmark of the “blind gut” model in early animal evolution. Here, we identify a pressure-sensitive aboral valve in *Nematostella vectensis* that operates independently of digestion. This valve dissipates elevated hydraulic pressure during morphogenesis, by expelling fluid through transient epidermal ruptures triggered by muscular ring opening. This unexpected function emerged from a comprehensive analysis of mesogleal basement membrane biogenesis. We show that the global dynamics of this extracellular matrix transduce muscular hydraulics to drive tissue rearrangement and stabilize shape, while localized FGFRb-dependent matrix remodeling establishes the aboral valve. By positioning the mesoglea as an integrator of biomechanics, tissue remodeling, and aboral valve function, these findings expand non-bilaterian openings beyond the digestive paradigm as a hydraulic regulator.

## Introduction

The emergence of multicellular animals required major innovations in tissue architecture and mechanical coordination. Central to this evolutionary transition was the extracellular matrix (ECM), a structural scaffold essential for shaping body plans and mediating morphogenetic signaling (*1–3*). In cnidarians—such as jellyfish, corals, and sea anemones—which represent one of the early branching metazoan lineages (*4*), the ECM is organized into a distinctive compartment known as the mesoglea, positioned between the ectoderm and endoderm within their diploblastic body plan (*5–7*). This fibrous interstitial matrix is bordered by basement membranes and has diversified across cnidarian lineages, ranging from thin sheets in sea anemones to voluminous matrices in jellyfish. The mesoglea offers critical insights into the primordial diversification of ECM-tissue functions.

Unlike vertebrates with rigid internal skeletons or arthropods with external exoskeletons, cnidarians rely on a hydrostatic system. In this system, the mesoglea acts as a dynamic elastic antagonist to muscle contractions, counterbalancing the pressure within their fluid-filled body cavity (*8*). This hydraulic mechanism generates and distributes internal pressure, enabling these seemingly simple animals to perform complex movements and maintain their body shape without hard structural elements (*9–12*). This dependence on internal pressure as a physiological driver underscores the need for precise regulatory mechanisms. Yet how this regulation is achieved and how it integrates with tissue architecture remains poorly understood. The prevailing “blind gut” model, in which the oral opening serves both ingestion and excretion, has long shaped models of early animal evolution (*13–15*). This anatomical paradigm distinguishes cnidarians from bilaterians, in which a through-gut evolved to compartmentalize digestive and excretory functions. However, whether this architectural simplicity accommodates specialized mechanisms for pressure regulation has not been explored.

Given the fundamental role of hydraulics in shaping cnidarian form and function (*8*, *12*, *16*), understanding how these forces are integrated with tissue architecture—particularly the mesoglea—is essential not only for uncovering the principles underlying their development and behavior, but also for informing bioinspired design in fields such as soft robotics (*13*, *14*). Despite its importance, the embryonic origin, assembly, and mechanical integration of the mesoglea are still unresolved. To address this, we leveraged the genetically tractable sea anemone *Nematostella vectensis* (*17*, *18*) to dissect mesogleal basement membrane biogenesis. During the larva-to-polyp transition, axial elongation is driven by muscular hydraulics and arrested by experimental depressurization (*16*). This period of morphogenesis coincides with extensive ECM remodeling, including the upregulation of matrix-modifying enzymes and basement membrane components like Collagen IV and Laminin (*19*). Together, these features make this developmental window an ideal context to investigate how mesogleal assembly interacts with hydraulic forces to orchestrate morphogenesis.

Here, we uncover the developmental origins and biomechanical integration of the mesogleal basement membrane during *Nematostella* morphogenesis. To achieve this, we combined genetic knock-ins, quantitative imaging, 3D electron microscopy, and molecular and biophysical perturbations. Unexpectedly, our analysis revises the classical view of cnidarian body architecture by revealing a pressure-sensitive aboral valve regulated by localized ECM remodeling and muscular control. During morphogenesis, this structure responds to elevated hydrostatic pressure within the body cavity by facilitating controlled fluid expulsion, thereby maintaining hydraulic pressure homeostasis.

## Results

### Embryonic origin of mesogleal basement membrane

To identify which embryonic tissue generates mesogleal basement membrane, we performed live imaging of the eGFP::ColIV knock-in (KI) line (*18*) during early development (Fig. 1A; Movie S1). At the blastula stage (∼12.5 hours post-fertilization, hpf), we observed that eGFP::ColIV was initially expressed in a polarized pattern marking the pre-endodermal plate prior to gastrulation. This expression intensified as these cells underwent invagination. To complement live imaging, we also examined fixed samples at multiple developmental stages, from the onset of gastrulation to the primary polyp (Fig. 1B). During gastrulation, eGFP::ColIV was primarily localized intracellularly in punctate structures, consistent with active synthesis. As development progressed, it transitioned to an extracellular localization upon completion of gastrulation, forming the basement membranes lining both germ layers (Fig. 1, B and C). Throughout this period, the mesoglea thickness remained relatively constant at approximately 2µm (Figure 1, C and D), while intracellular eGFP::ColIV was consistently restricted to the endoderm/gastrodermis (Fig. 1C). Within this tissue, it displayed a distinct subcellular distribution, with large apical puncta and smaller basal puncta. Given their proximity to the forming ECM, these basal puncta likely represent sites of Collagen IV secretion.

**Figure 1.**
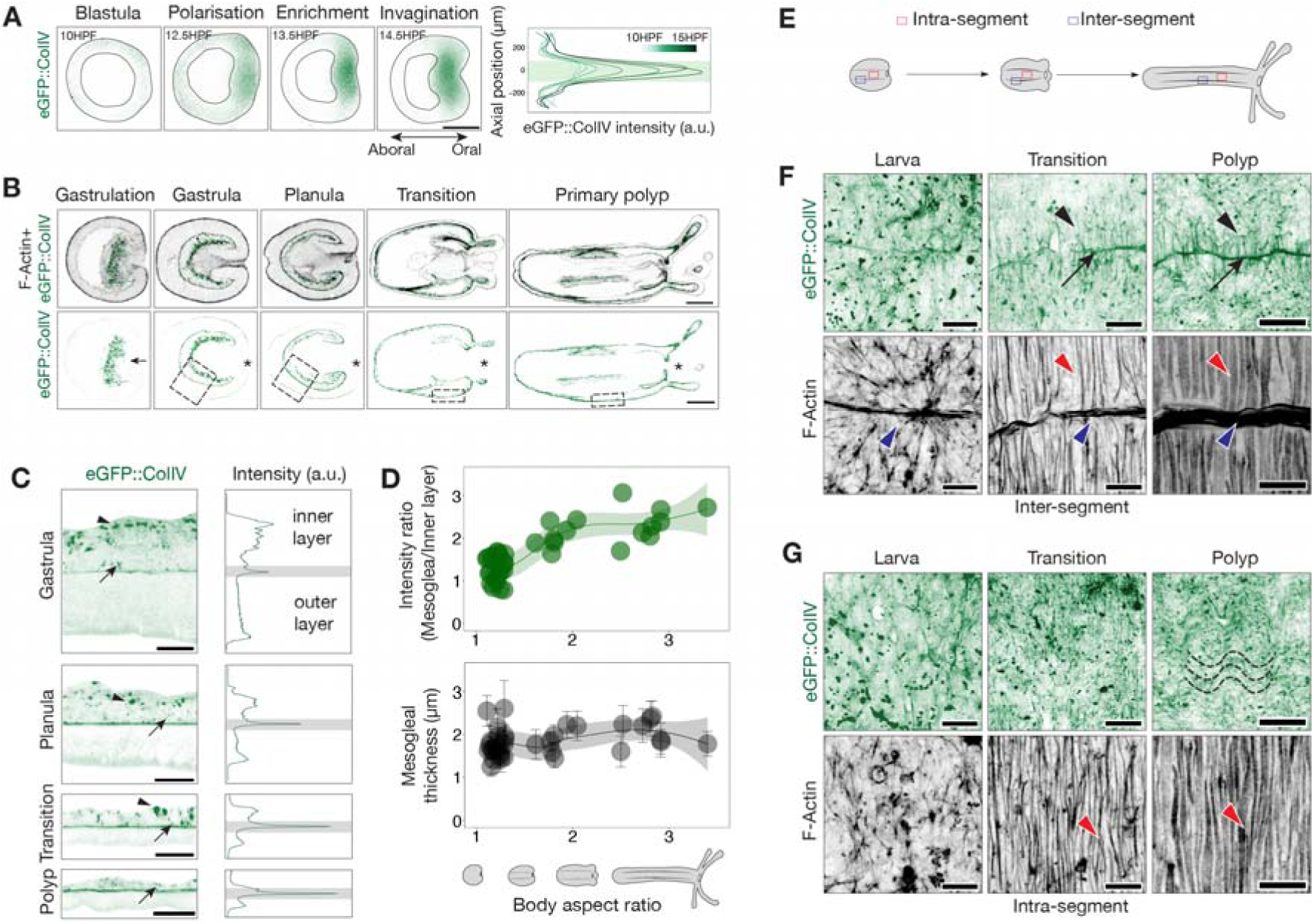
Developmental dynamics of mesogleal basement membrane. (A) Lightsheet imaging of a live eGFP::ColIV embryo. Left: Optical cross-section of a developing blastula showing endogenous Collagen IV expression (green) before the onset of gastrulation. Right: Quantification of eGFP::ColIV intensity along the oral–aboral axis (0: oral pole). Scale bar: 100 µm. (B) Confocal cross-sections of fixed eGFP::ColIV-positive animals from gastrulation to the primary polyp stage. F-actin is stained in gray. Scale bar: 100 µm. (C) Magnified insets from B with corresponding average intensity profiles along the xy-axis. The central plane represents the developing mesoglea.; upper and lower regions correspond to endoderm and ectoderm, respectively. The intensity of the extracellular eGFP::ColIV is highlighted in gray. Apical and basal eGFP::ColIV puncta are indicated by arrowhead and arrow, respectively. Scale bar: 100 µm. (D) Ratio of extracellular (mesoglea) to intracellular (inner layer) *eGFP::ColIV* intensity as a function of body aspect ratio, *n =* 50. (E) Schematic illustrating the spatial localization of intra-segmental and inter-segmental regions during development. (F) Maximum intensity Z-projection of inter-segmental regions in *eGFP::ColIV*-positive animals (green) with F-actin staining (gray). Black arrowheads indicate short lateral Collagen IV bridges; black arrows mark Collagen IV accumulation at inter-segmental boundaries. Red and blue arrowheads indicate circular and parietal muscles, respectively. Scale bar: 25 µm. (G) Maximum intensity Z-projection of intra-segmental regions in *eGFP::ColIV*-positive animals (green) with F-actin staining (gray). Dashed lines indicate the wavy pattern of Collagen IV network. Scale bar: 25 µm.

To track the temporal dynamics of Collagen IV deposition, we quantified both intracellular and extracellular eGFP::ColIV fluorescence intensities throughout development (Fig. 1, C and D; fig. S1A). These measurements were correlated with body aspect ratio (A/R; length divided by width), as a proxy for developmental progress during axial elongation. As eGFP::ColIV was uniformly distributed throughout the central body region (fig. S1A), we calculated the average local fluorescence intensities at each stage. We observed that extracellular eGFP::ColIV levels increased from gastrula until mid-planula, then plateaued during later development (Fig. 1D). This pattern was recapitulated by immunostaining for Collagen IV (*20*) and laminin (*19*) (fig. S1B), although Collagen IV stabilized earlier than laminin, suggesting a sequential assembly of basement membrane components. Collectively, these results demonstrate that the endoderm is the main and continuous source of developmentally regulated Collagen IV production.

### Developmental patterning of mesogleal basement membrane

After characterizing Collagen IV production across development, we next investigated how it is spatially organized, along with Laminin, to form a functional basement membrane. In gastrula, Collagen IV displayed a diffuse distribution, while Laminin was not yet detectable (fig. S2, A and B). In early larval stages, the diffuse pattern of Collagen IV began to resolve into a more structured arrangement (fig. S2B), while Laminin incorporated broadly across the tissue and exhibited early signs of spatial organization. As development progressed, localized enrichment of both Collagen IV and Laminin emerged at sites of developing endodermal folds, marking the segment boundaries of future gastrodermal mesenteric folds (Fig. 1, E and F; fig. S2). These segment boundaries (inter-segments) continued to accumulate Collagen IV and Laminin, and new structures appeared—short lateral bridges spanning adjacent segments that were marked by Collagen IV, but not Laminin (Fig. 1F; fig. S2B). The spatial density of these bridges increased progressively throughout development. Within the body wall segments (intra-segments), both Collagen IV and Laminin adopted an undulating arrangement that became increasingly pronounced in polyps (Fig. 1G; fig S2B). These spatial patterns were consistently observed in both the eGFP::ColIV KI line and animals stained for Collagen IV and Laminin.

To examine how this structural organization emerges, we used photoconversion experiments with the Dendra2::ColIV KI line (*21*) to track ECM remodeling over time. In larvae, we photoconverted discrete lateral patches of Dendra2::ColIV (magenta) in intra-segment regions (fig. S2, C and D). These patches realigned into an axial undulating configuration and incorporated newly synthesized Dendra2::ColIV (green) (Fig. 1I), indicating that pre-existing Collagen IV is continuously being remodeled as part of a dynamically evolving network. We also analyzed Collagen IV dynamics in inter-segment regions (fig. S2, E and F) to determine whether lateral bridges formed through sequential addition or by intercalation among existing ones. If formation occurred via sequential addition, photoconverted (magenta) and newly synthesized (green) bridges would remain spatially distinct. Instead, we observed extensive mixing between old and new bridges, supporting an intercalation model in which new bridges integrate between pre-existing ones (Fig. 1K). These results reveal the remodeling of the developing mesogleal basement membrane which progressively transitions from a diffuse, unstructured matrix to a spatially organized scaffold during morphogenesis.

### Endodermal morphogenesis drives basement membrane organization

To investigate the cellular processes underlying basement membrane organization, we examined the relationship between endodermal morphogenesis and ECM architecture. During development, the endoderm undergoes both tissue folding and differentiation into a spatially patterned musculature composed of circular and longitudinal muscles (*22*). These morphogenetic processes may influence basement membrane organization through mechanical interactions and/or localized ECM production. To simultaneously visualize Collagen IV organization and muscle development, we combined the eGFP::ColIV KI line with F-actin staining (Fig. 1, F and G). In early larvae, Collagen IV intensity at endodermal folds progressively increased in parallel with the differentiation of parietal longitudinal muscles into thick bundles (Fig. 1F). At the same time, the emergence of short lateral bridges coincided with the formation of circular muscles (Fig. 1F). As development proceeded, both the lateral bridges and the intra-segmental undulating Collagen IV pattern became more pronounced, paralleling the maturation of circular muscles (Fig. 1G). To test whether endodermal morphogenesis influences basement membrane architecture, we performed targeted perturbations of key developmental regulators. BMP knockdown (KD), which disrupts endodermal folding and abolishes longitudinal muscle formation (*23*, *24*), resulted in a complete loss of Collagen IV and Laminin at segment boundaries, including the lateral bridges (fig. S2G). Tbx20 KD, which impairs muscle patterning (*16*), also resulted in a dramatic disorganization of both Collagen IV and Laminin (fig. S2G). These findings indicate that the architecture of the mesogleal basement membrane is actively shaped by endodermal morphogenetic processes.

### ECM modulators alter basement membrane composition

Having characterized the biogenesis of the mesogleal basement membrane, we next investigated its functional role during development. We initially performed Collagen IV KD experiments using shRNA (*25*) in the eGFP::ColIV knock-in (KI) line. In Collagen IV KD embryos, eGFP fluorescence was completely abolished, confirming efficient KD (fig. S3A). These embryos exhibited severe tissue disorganization in both the ectoderm and endoderm, including a failure of adhesion between the two germ layers (fig. S3B). Given the essential role of Collagen IV in early embryogenesis, we shifted our focus to perturbing basement membrane remodeling during post-embryonic development. To this end, we used two pharmacological inhibitors targeting distinct steps in ECM regulation (fig. S3C): GM6001, a broad-spectrum matrix metalloproteinase (MMP) inhibitor (*21*), and 2,2’-bipyridine (BPY), an inhibitor of prolyl-4-hydroxylase (P4HA) (*26*). GM6001 blocks ECM degradation, leading to accumulation of ECM components, whereas BPY impairs collagen hydroxylation, preventing proper polymerization and integration of Collagen IV into the basement membrane. These treatments were expected to alter ECM composition and, in turn, modulate the biomechanical properties of the mesoglea.

Pharmacological treatments were applied to 3-day-old planula larvae and maintained for three days during the larva-to-polyp transition. GM6001 treatment led to Collagen IV accumulation and mesoglea thickening (Fig. 2, A and B), while Laminin levels remained largely unchanged. Despite the increased Collagen IV deposition, the overall intra- and inter-segmental organization of Collagen IV and Laminin was largely preserved (Fig. 2A). In contrast, BPY-treated larvae showed a substantial reduction in Collagen IV level, whereas Laminin level was relatively unaffected (Fig. 2A). The basement membrane in BPY-treated animals appeared loosely organized and highly irregular, with increased variability in mesoglea thickness compared to controls (Fig. 2B). Together, these results demonstrate that GM6001 and BPY exert distinct and opposing effects on Collagen IV levels, enabling us to experimentally perturb the steady state of Collagen IV accumulation that normally stabilizes after mid-planula stage.

**Figure 2.**
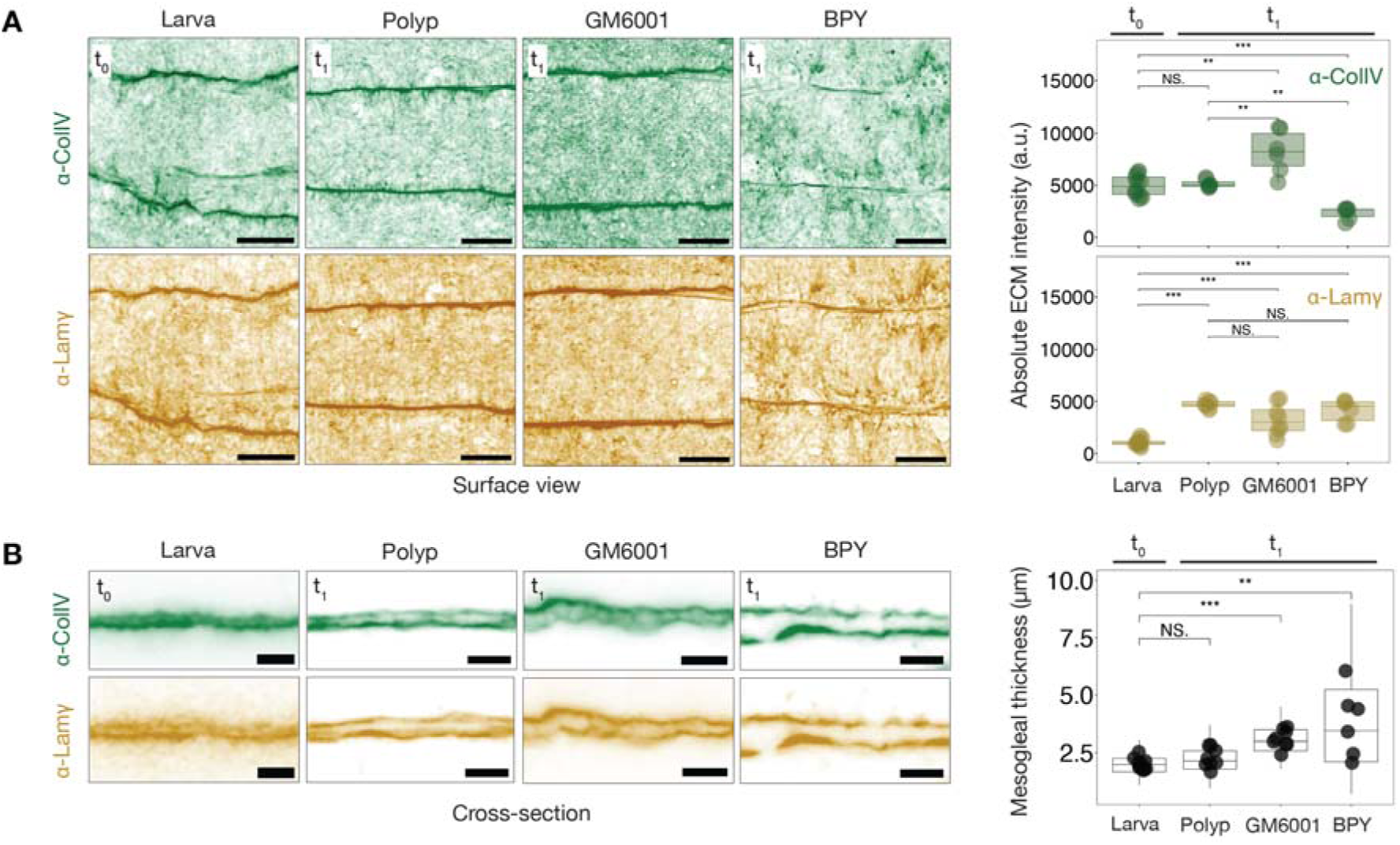
Perturbations of basement membrane composition. **(A)** (Left) Maximum intensity projections showing Collagen IV and Laminin immunostaining in untreated (t_0_) and treated (t_1_, 3 days post-treatment) larvae under the indicated conditions. Scale bar: 20 µm. (Right) Quantification of average fluorescence intensity for Collagen IV and Laminin. *n =* 10 larvae; *n* = 8 DMSO-treated polyps; *n* = 8 GM6001-treated animals; *n* = 8 BPY-treated animals. **(B)** (Left) Cross-sectional views of the basement membrane stained for Collagen IV and Laminin under the indicated conditions. Scale bar: 20 µm. (Right) Quantification of mesoglea thickness across conditions. *n* = 12 larvae; *n* = 12 DMSO-treated polyps; *n* = 12 GM6001-treated; *n* = 6 BPY-treated animals.

### Mesoglea integrity controls axial elongation and hydraulic homeostasis

We next used quantitative live imaging to determine how GM6001 and BPY treatments influence larva-to-polyp morphogenesis—a process that requires coordinated changes in body shape and size driven by muscle-generated fluid pressure (16) (Fig. 3A; Movie S2). Both treatments arrested axial elongation midway through the developmental trajectory (Fig. 3A; fig. S4A; Movie S2), indicating that intact ECM remodeling is essential for successful morphogenesis. However, their trajectories following arrest diverged markedly. GM6001-treated animals remained morphologically stable after arrest, whereas BPY-treated animals exhibited a progressive reversal of elongation, characterized by reductions in both shape and size (Fig. 3A; Movie S2). Strikingly, this regression in BPY-treated animals was accompanied by abrupt leakage from the aboral pole, during which internal cavity contents were expelled into the surrounding medium (highlighted by dashed lines in Fig. 3B; Movie S3). This aboral leakage, presumably driven by elevated internal cavity pressure (16), led to a rapid loss of cavity volume and a corresponding decrease in body size (Fig. 3, A and B). To determine whether this phenomenon was specific to BPY treatment, we re-examined elongation dynamics in GM6001-treated and control animals. No leakage events were observed in GM6001-treated animals (Movie S2), while control animals exhibited occasional aboral leakage near the end of elongation (Fig. 3B; Movie S3), indicating that pressure release at the aboral pole may occur naturally but is normally restricted to later developmental stages. These observations suggest the presence of a previously unrecognized pressure-responsive zone at the aboral pole that facilitates fluid release during morphogenesis (Fig. 3C). Under normal conditions, this region resists leakage during early development, but its structural integrity is compromised when ECM composition is disrupted.

**Figure 3.**
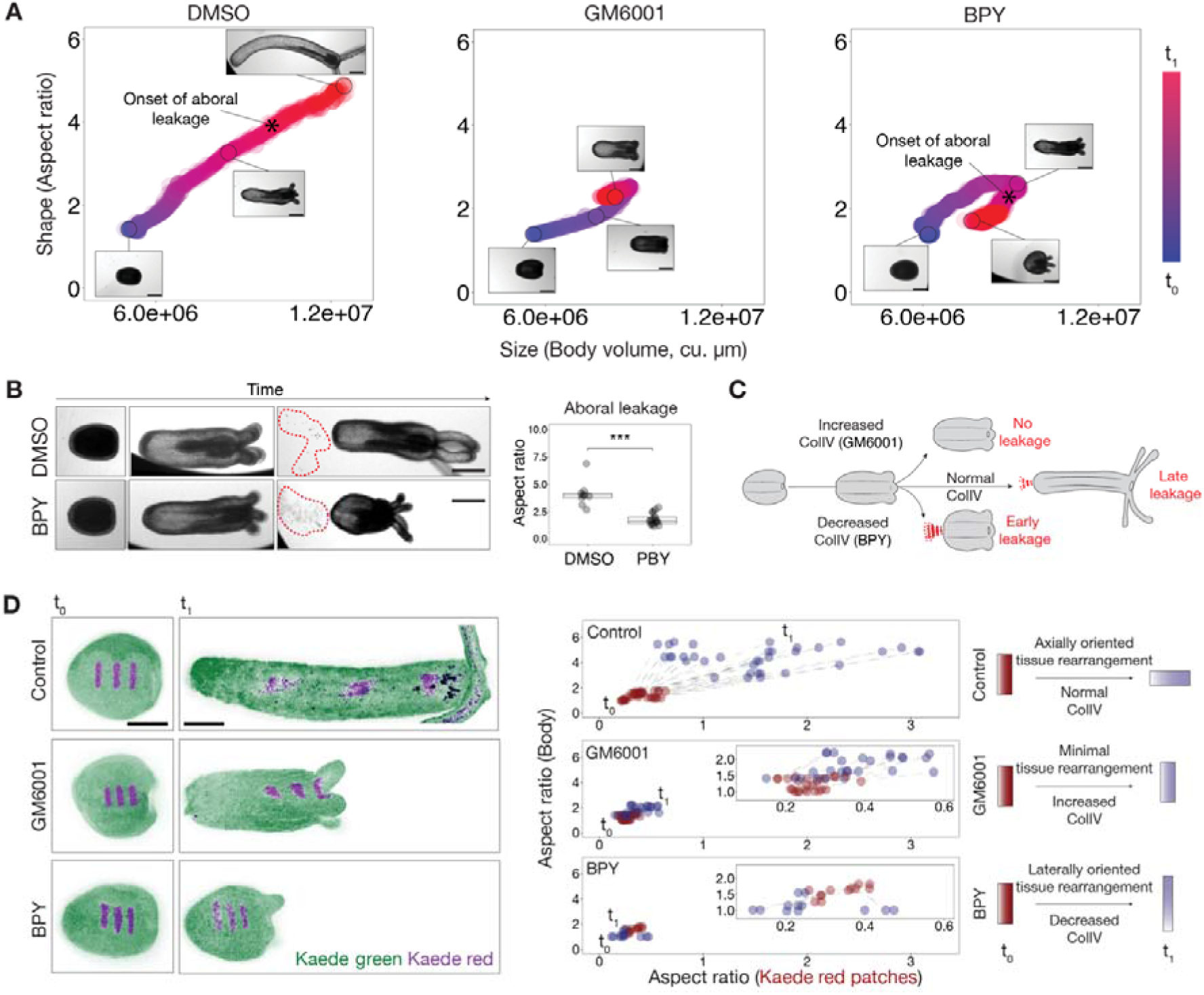
ECM modulation disrupts tissue rearrangement and hydraulics. **(A)** Morphospace analysis showing elongation trajectories based on changes in body volume (size) and aspect ratio (shape) from t_0_ to t_1_ (3-day interval) under the indicated conditions. Onset of aboral leakage is indicated in the developmental trajectory. *n* = 20 DMSO-treated; *n* = 20 GM6001-treated; *n* = 20 BPY-treated animals. **(B)** (Left) Aboral leakage phenotypes in control and BPY-treated animals. Note the cloud of cellular debris expelled from the oral pole. Scale bar: 100µm. (Right) Quantification of the onset of aboral leakage as a function of body aspect ratio. *n* = 10 DMSO-treated; *n* = 10 BPY-treated animals. **(C)** Schematic representation of aboral leakage observed across experimental conditions. **(D)** (Left) Maximum intensity projection images of photoconverted tissue patches in larvae and corresponding topological changes at t_1_. Green: non-photoconverted; magenta: photoconverted. Scale bar: 100 µm. (Right) Quantification of patch elongation (length/width aspect ratio) versus body column aspect ratio at each time point across experimental conditions. (Right) Schematic summarizing trends in patch shape dynamics across treatments.

More broadly, these experiments indicate that a finely tuned mesoglea composition is critical not only for sustaining body elongation and maintaining overall shape but also for preserving internal hydraulic homeostasis throughout morphogenesis. Inhibition of matrix metalloproteinases by GM6001 reduces ECM turnover and promotes collagen accumulation, reinforcing the mesoglea and enabling tissues to withstand internal pressure—though at the cost of preventing tissue remodeling and arresting elongation. In contrast, BPY treatment destabilizes and weakens the mesoglea, rendering the tissue unable to sustain hydraulic stress. This effect results in both a failure to elongate and premature aboral leakage, culminating in a progressive loss of shape stability and hydraulic homeostasis. In the following sections, we dissect the mechanisms by which the mesoglea (*i*) supports body elongation and (*ii*) modulates aboral leakage during morphogenesis.

### Global ECM modulation impairs axial tissue rearrangement

Since body elongation during the larva-to-polyp transition depends primarily on tissue remodeling rather than cell proliferation (*16*), we investigated how GM6001 and BPY treatments affect the underlying cellular mechanisms. In previous work, we showed that elongation proceeds through two distinct phases: an initial stage characterized by changes in cell shape that thin the body wall and enable surface expansion, followed by a phase of oriented tissue rearrangement that drives directional elongation along the oral–aboral axis (*16*). In control animals, tissue thinning accounted for early body shape changes, whereas continued elongation was achieved through axial tissue reorganization (Fig. 3D; fig S4B).

Because GM6001- and BPY-treated animals arrested elongation at an intermediate stage and phenocopied the effects of muscle anesthetics (*16*) (Fig. 3A; fig S4B), we hypothesized that ECM perturbation disrupts muscle-driven tissue rearrangement without impairing body wall thinning. Consistent with this, drug-treated animals exhibited degrees of body wall thinning comparable to controls (fig S4B), while photoconversion experiments using Kaede-labeled larval tissue revealed striking defects in axial tissue rearrangement (Fig. 3D). In GM6001-treated animals, photoconverted patches showed minimal elongation along the body axis, indicating limited rearrangement (Fig. 3D). In contrast, BPY-treated animals exhibited lateral expansion of photoconverted patches rather than axial elongation, consistent with their reversal of body elongation and rounded morphology (Fig. 3D). These results suggest that ECM modulation alters mesoglea organization in ways that specifically interfere with axial tissue rearrangement driven by muscular hydraulics.

### Localized mesoglea remodeling supports a pressure-sensitive aboral valve

In addition to its global role in tissue remodeling, the mesoglea may contribute to region-specific functions during morphogenesis. Early aboral leakage in BPY-treated animals and its delayed onset in developing wild-type polyps suggest a localized difference in tissue behavior at the aboral pole. These observations raise the possibility of spatial heterogeneity in mesoglea properties. To explore this possibility, we examined the structural organization of the aboral pole in greater detail. We first analyzed the distribution of Collagen IV and Laminin in primary polyps (Fig. 4A). At the aboral pole, Collagen IV displayed a localized reduction, forming a discrete ∼5□µm-wide region with diminished signal. In contrast, Laminin remained broadly distributed but showed local thickening in the same area, consistent with spatially restricted basement membrane remodeling. To further examine tissue architecture, we performed focused ion beam scanning electron microscopy (FIB-SEM) on a primary polyp, acquiring a volume of 3500 µm³ at nanometer-scale resolution, which we subsequently segmented to characterize tissue organization (Fig. 4B, Movie S4). Despite the occurrence of fluid leakage, we found no evidence of a persistent discontinuity in the epidermal surface, ruling out the presence of a continuously open pore. Instead, we identified a rosette-like arrangement at the basal epidermis, where neighboring cells had lost direct contact, forming a structured extracellular space (Fig. 4B, Movie S4). Within this space, protruding gastrodermal cells were observed, indicating a specialized aboral tissue architecture.

**Figure 4.**
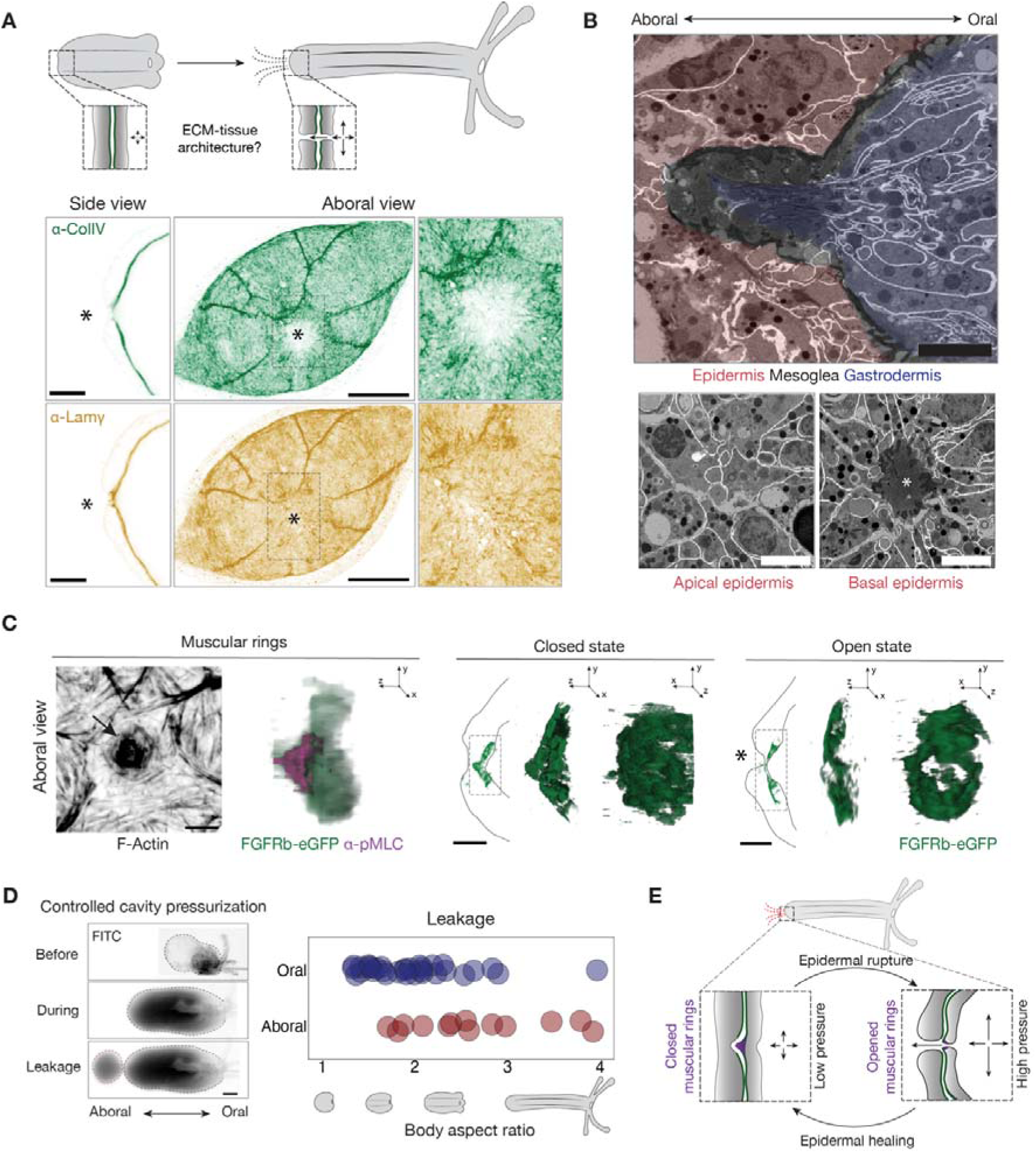
Localized ECM remodeling defines an aboral valve. **(A)** (Top) Schematic depicting the tissue and ECM remodeling events required for aboral leakage during the larva-to-polyp transition. (Bottom) Cross-sectional (side view) and maximum intensity projection (aboral view) images showing Collagen IV and Laminin immunostaining in polyps. Asterisk indicates the site of ECM remodeling. Scale bar: 20 µm. **(B)** (Top) Cross-sectional view of Segmented FIB-SEM data revealing the architecture of the aboral pole. Scale bar: 5µm. (Bottom) Aboral view of the FIB-SEM dataset highlighting epidermal architecture. Asterisk marks the basal gap in the epidermis. Scale bar: 10 µm. **(C)** (Left) F-actin and phospho-Myosin Light Chain (pMLC) staining of muscular rings reveals its contractile organization (see arrow). Scale bar: 20 µm (top), 10µm (bottom). (Right) Cross-sectional view and 3D rendering showing the dual conformational states of muscular rings labeled with *FGFRb-eGFP*. Asterisk indicates epidermal discontinuity. Scale bar: 20 µm. **(D)** (Left) Cavity inflation assay in a primary polyp showing three representative time points: before inflation, during inflation, and at the point of leakage. Scale bar: 50 µm. (Right) Quantification of oral versus aboral leakage events as a function of body aspect ratio across developmental stages. **(E)** Schematic model summarizing the mechanism of aboral valve function.

To determine whether this aboral tissue architecture is consistently present across individuals, we imaged FGFRb-eGFP transgenic polyps (*27*), in which a cluster of gastrodermal cells at the aboral pole is fluorescently labeled (*21*). These clusters adopted two distinct configurations: a funnel-like structure with a continuous surface or a flattened structure featuring a central gap (Fig. 4C; Movie S5), suggesting that the gastrodermal interface cycles between closed and open states. Additionally, FGFRb-eGFP–positive clusters formed concentric muscle rings characterized by enrichment of F-actin and phosphorylated myosin light chain (p-MLC) (Fig. 4C). This enrichment was particularly prominent at the tip of the funnel structure, suggesting that muscles actively control the configuration of this anatomical feature. To confirm that this structure functions as a pressure-responsive valve, we artificially inflated the body cavity by introducing a dye-filled capillary through the mouth, connected to a pressure pump (Fig. 4D, Movie S6). Upon pressurization, dye was consistently expelled through the aboral pole rather than the mouth, showing that this anatomical feature serves as a directional release valve activated by elevated internal pressure (Fig. 4D). Interestingly, the epidermis above the muscular rings alternated between continuous and disrupted states (Fig. 4C), suggesting that pressure release transiently induces epithelial rupture.

To test whether this rupture triggers a wound response, we examined ERK signaling, a conserved marker of epithelial injury across metazoans (*28*). In control polyps, phosphorylated ERK (pERK) was variably activated in the aboral epidermis, with signal intensity ranging from low to high across individuals (fig. S5). In contrast, physically injured polyps exhibited robust pERK activation in both the aboral epidermis and gastrodermis. These results suggest that pERK activation in uninjured polyps reflects transient, pressure-induced epithelial wound.

Collectively, the data support the existence of an aboral valve that mediates internal pressure release through cycles of epithelial disruption and repair. Acting as a “physiological wound,” this dynamic structure enables rapid decompression when the oral opening is sealed, with its activity regulated by muscle control and localized ECM remodeling (Fig. 4E).

### FGFRb-dependent aboral valve morphogenesis

To investigate how the aboral valve forms during development, we examined the organization of Collagen IV and Laminin at the aboral pole across developmental stages. In embryos, Collagen IV was uniformly distributed throughout the mesoglea, with Laminin undetectable (fig. S2A). During the larval stage (fig. 6A), Collagen IV became progressively depleted across a broad region of the aboral mesoglea, forming a distinct depletion zone that contracted in size as development proceeded. Laminin was also reduced in this region during the larval period. However, during the larva-to-polyp transition (fig. 6A), Laminin levels were restored and became specifically enriched within the Collagen IV-depleted zone. These spatially divergent remodeling patterns were disrupted in GM6001- and BPY-treated animals (fig. 6B), suggesting that mesoglea weakening at the aboral pole is a temporally coordinated process.

Despite these early signs of ECM remodeling, pressurized larvae consistently expelled fluid through the mouth (Fig. 4D; Movie S6). In contrast, animals undergoing the larva-to-polyp transition exhibited variable fluid release—either orally or aborally—until they reached a more elongated polyp stage, when aboral release became predominant (Fig. 4D; Movie S6). This shift likely reflects the progressive maturation of contractile muscular rings at the aboral pole. Supporting this, FGFRb-eGFP–positive concentric muscle rings emerged only during late transition stages and coincided with strong enrichment of p-MLC at the aboral pole, a hallmark of an actively contractile structure (fig. 6A).

To test whether FGFRb signaling is required for aboral valve morphogenesis, we analyzed FGFRb mutant animals (*27*). Unlike their siblings, FGFRb mutants failed to exhibit localized Collagen IV depletion at the aboral pole (Fig. 5A), and their musculature was highly disorganized, lacking the characteristic p-MLC enrichment observed in controls (Fig. 5B). These structural defects suggest a failure to form a functional aboral valve. To test this, we artificially inflated the body cavities of FGFRb mutants. While control animals consistently expelled dye through the aboral pole, mutants released fluid exclusively through the mouth, with only a single exception (Fig. 5C; Movie S7). Furthermore, mutants required significantly higher internal pressure to trigger fluid expulsion compared to controls, confirming that aboral release is mechanically impaired in the absence of FGFRb signaling.

**Figure 5.**
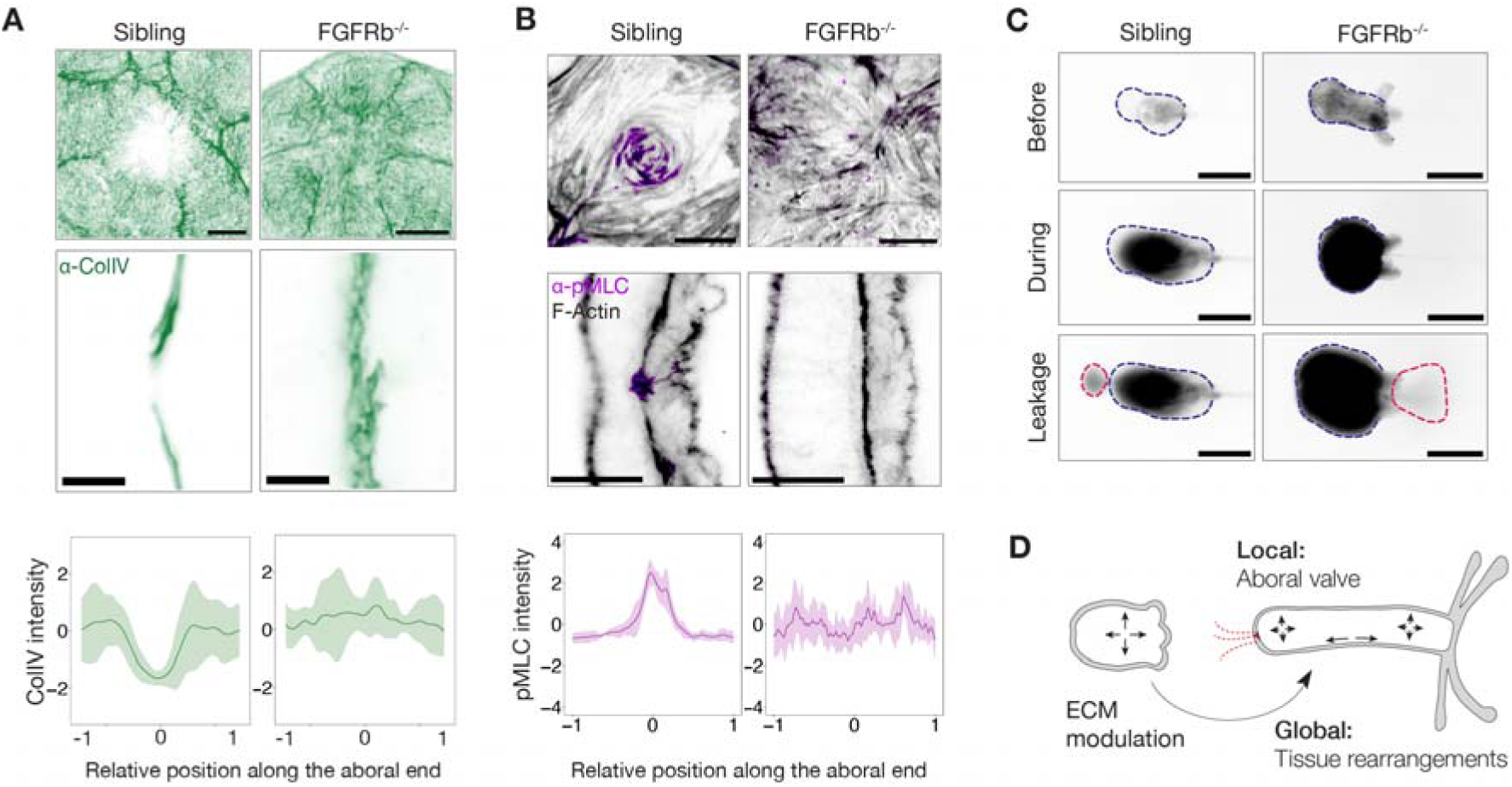
Developmental basis of aboral valve formation. **(A)** (Top) Maximum intensity projections (aboral view) showing Collagen IV distribution in *FGFRb* knockout (*FGFRb-/-*) mutants and representative siblings. (Middle) Cross-sectional views of the aboral pole in the respective genotypes. Scale bar: 10µm. (Bottom) Quantification of Collagen IV spatial distribution in *FGFRb-/-* mutants versus siblings. **(B)** (Top) Maximum intensity projections (aboral view) showing F-actin and pMLC staining in *FGFRb-/-* mutants and representative siblings. (Middle) Cross-sectional views of the aboral pole in each genotype. Scale bar: 20µm. (Bottom) Quantification of pMLC signal distribution in *FGFRb-/-* mutants versus siblings. **(C)** Cavity inflation assay in *FGFRb-/-* mutants and siblings. Aboral leakage is observed in 1 out of 9 *FGFRb-/-* mutants, compared to 7 out of 9 siblings. Scale bar: 200 µm. **(D)** Schematic model illustrating the processes dependent on local and global mesoglea remodeling.

Together, these findings demonstrate that FGFRb signaling is essential for coordinating the localized ECM remodeling and muscle ring assembly required to form a functional, pressure-sensitive aboral valve.

## Discussions

Our study reveals the dual roles of the mesogleal basement membrane in cnidarian morphogenesis, operating at both global and local scales (Fig. 5D). At the global level, we demonstrate that the dynamic mesoglea architecture orchestrates tissue remodeling and axial elongation. At the local level, we identify a specialized mesogleal domain at the aboral pole that undergoes targeted remodeling to create a pressure-sensitive valve, a previously unrecognized function that revises the traditional view of cnidarian body architecture.

We demonstrate that Collagen IV is expressed precociously in presumptive endoderm cells prior to gastrulation, which likely stabilizes initial contacts between invaginating endodermal filopodia and the basal blastoderm surface (*29*, *30*). This early expression facilitates bilayer formation and enables subsequent morphogenetic events. Throughout development, endoderm-derived Collagen IV forms a regulated ECM that dynamically interacts with muscular hydraulics (*18*), essential for directional tissue rearrangements during axial elongation. Excess Collagen IV prematurely restricts tissue rearrangement, halting elongation, whereas reduced collagen destabilizes morphology through misaligned remodeling. These results suggest that mesoglea mechanics must be finely tuned, allowing plastic deformation (*31*, *32*) without compromising structural integrity. In parallel, the enrichment of Wnt/PCP signaling components in mesoglea proteomes (*19*) suggests integration of mechano-chemical signaling in larva-polyp morphogenesis, an avenue for future investigation.

Although scattered reports have noted possible secondary openings in cnidarians (*33–35*), such claims have historically been dismissed due to the fragility of cnidarian tissues. Here, we provide definitive structural and functional evidence for a dynamic aboral valve. This site features localized ECM remodeling and contractile muscle rings, and opens transiently under internal pressure through controlled epithelial rupture and wound signaling activation. Crucially, this is not a through-gut. Unlike the permanent anal pore in ctenophores (*15*), which support unidirectional digestion, the aboral opening in *Nematostella* is transient, muscle-controlled, and pressure-responsive. It functions not in waste expulsion, but as a biomechanical safety valve to offload internal pressure. While pressure and fluid could also be released through the oral opening, this aboral valve likely serves as a “backup exit” when oral release is obstructed, such as during pharyngeal compression (fig. S7). This dual-exit system introduces mechanical redundancy, ensuring robust physiological control of internal pressure.

Importantly, this pressure-release mechanism may confer adaptive advantages in the brackish coastal habitats of *Nematostella*, where rapid salinity fluctuations frequently occur (*36*). A stress-induced epithelial rupture also occurs at the oral pole in Hydra during feeding (*37*, *38*), suggesting that stress-responsive epithelial discontinuities may be an ancient feature of cnidarian biology. Together, our findings redefine the concept of body openings in early branching animals. Rather than viewing epithelial perforations solely through the lens of unidirectional gut evolution, we propose that transient, muscle-regulated rupture zones may have served ancestral roles in pressure regulation. More broadly, these findings illuminate how coordinated global and local ECM dynamics collectively shape organismal form and function in one of the earliest-diverging animal lineages with organized tissue layers.

## Acknowledgments

We thank Kresimir Crnokic for his support in animal husbandry. We thank the Ikmi lab members for their comments on the work. We also thank Muzamil Majid Khan for the discussion about the ECM and for sharing ECM-related reagents, as well as the feedback on the manuscript.

## Author Contributions

S.B. and A.I. conceived the idea for this project and designed the experiments. S.B. performed most of the experiments. A.P. designed the KI lines. P.S. performed immunostaining and drug treatments. F.G. performed lightsheet imaging. G.B. T.H. and P.R. processed the sample and performed the experiments for FIB-SEM. J.H. and A.K. segmented the FIB-SEM data. S.O. generated the Laminin antibody. S.B. and A.I. analyzed the data. S.B. generated all figures. A.I. drafted the manuscript with inputs from all authors.

## Data and materials availability

All data needed to evaluate the conclusions are present in the paper and the supplementary materials. Transgenic lines are available upon request.

## Materials and Methods

### Animal husbandry and Spawning

Adult *Nematostella vectensis* were cultured in 12 ppt artificial seawater (ASW; Instant Ocean sea salt) at 17°C under dark conditions. To induce spawning, animals were placed in a white light incubator for 6–8 hours overnight, with the temperature increased to approximately 28°C (*1*). Spawning typically occurred within 3–4 hours following a cold water change (17°C). Collected eggs were de-jellied by incubating them for 9 minutes in a 4% cysteine solution (Sigma, 168149) prepared in ASW, then rinsed three times with fresh ASW prior to fertilization.

### Transgenic and mutant lines

The *eGFP::ColIV* and *Dendra2::ColIV* knock-in lines used in this study were previously generated (*2*, *3*). The *FGFRb-eGFP* reporter and *FGFRb KO* lines were described earlier (*4*). Transgenic embryos were obtained by crossing transgenic males with wild-type females.

### Live imaging of early embryos

Live confocal imaging was performed on a Leica SP8 CSU confocal microscope equipped with a 20× objective, with embryos mounted non-confined in 0.22 µm-filtered 12 ppt artificial sea water on MatTek round glass-bottom dishes (#P35G-1.5-14-C) and imaged overnight.

Live light-sheet imaging was performed on a Luxendo MuVi-SPIM using a Nikon CFI Plan Fluor 10×/0.30 NA illumination objective and an Olympus XLUMPLFLN 20×/1.00 NA detection objective, both immersed in 0.22 µm-filtered 12 ppt sea water and maintained at 23 °C; fluorescence was excited at 488 nm (∼1.6 mW) with a 3 µm beam width, 50 ms exposure, and captured on a Hamamatsu C11440-22C camera. Embryos were mounted non-confined on a 1% agarose in filtered 12 ppt sea water bedding in 100 µL glass capillaries trimmed for the sample holder, enclosed in FEP tubing. Prior to mounting the embryos onto the bedding, the agarosesolidified in the glass capillaries at 4 °C for 10 min, to then fill the FEP tube with sea water to eliminate bubbles. Time-lapse Z-stacks (1 µm steps) were recorded every 5 min for 12.5–40 h, acquiring four orthogonal views per time point (two detection arms plus a 90° rotation of each).

### Acquifer microscope

Live imaging of the larva-to-polyp transition was conducted using an Acquifer screening microscope (*5*). Larvae (3 dpf) were individually placed into wells of a 384-well plate (Corning, 3540), each containing 25□µL of 12 ppt artificial seawater (ASW) with 0.1% DMSO or pharmacological inhibitors. Imaging was performed continuously every 5 minutes over three days at 27□°C, using a 4× objective and brightfield illumination set to 20% intensity. Image analysis was performed as previously described (*5*).

### Immunostaining

Animals were first anesthetized in 7% magnesium chloride prior to fixation. Fixation was performed at room temperature for 1 hour using either 4% paraformaldehyde in PBS (Electron Microscopy Sciences, E15710) for transgenic line samples or Lavdovsky’s fixative (3.7% formaldehyde, 50% ethanol, 4% acetic acid) for immunostaining of basement membrane components. Fixed samples were treated for 20 minutes in 10% DMSO (Thermo Fisher, 85190) in PBS, followed by rinses in PBS containing 0.2% Triton X-100 (Sigma, T8787) (PTx 0.2%). Blocking was carried out for 1 hour in PTx 0.1% supplemented with 0.1% DMSO, 1% BSA (Sigma, A2153), and 5% goat serum (Sigma, 8182G9023). Samples were then incubated overnight at 4□°C with primary antibodies diluted in blocking buffer (Table 1). After thorough washing in PTx 0.1%, Alexa Fluor–conjugated secondary antibodies (Thermo Fisher, 1:500) were applied in the same buffer and incubated overnight at 4□°C. For additional labeling, F-actin was stained with Alexa Fluor–conjugated phalloidin (Thermo Fisher, 1:100), and nuclei were counterstained with Hoechst 34580 (Sigma, 63493, 1:1000), both in PTx 0.1% overnight at 4□°C. Finally, all samples were washed in PTx 0.1% and mounted in Vectashield Plus (Vector Laboratories) for confocal microscopy.

**Table 1:** Primary antibodies used for immunostaining and their working dilutions.

| Antibody | Source | Dilution |
| --- | --- | --- |
| anti-phospho-Myosin Light Chain | Cell Signaling Technology #3671S | 1:50 |
| anti-eGFP | MBL #598S | 1:500 |
| anti-eGFP | Abcam #ab1218 | 1:500 |
| anti-Laminin-γ | Bergheim et al., 2025 | 1:400 |
| anti-ColIV - JK2 | Gift from Haruko Tomono | 1:400 |
| anti-phospho-p44/42 MAPK (Erk1/2)(Thr202/Tyr204) | Cell Signaling Technology #4370 | 1:250 |

### Direct visualisation of reporter lines

Animals expressing transgenes or KI constructs were anesthetized in 7% magnesium chloride before fixation. Fixation was carried out for one hour at room temperature using 4% paraformaldehyde (EMS, E15710) in PBS. Afterwards, the samples were washed four times for 5 minutes each with 1X PBS. Additional staining for F-actin and nuclei was performed using phalloidin Alexa Fluor (Thermo Fisher, 1:100) and Hoechst 34580 (Sigma, 63493, 1:1000), respectively, in 1X PBS for 6 hours at 4°C. The samples were protected from light post-fixation. Finally, the samples were directly mounted in Vectashield Plus for confocal imaging.

### Confocal imaging

Samples were imaged using either a Zeiss LSM880 AiryFast or Zeiss LSM980 AiryFast microscope. For lower-resolution acquisition, a Plan-Apochromat 20×/0.8 M27 air objective or an LD-LCI Plan-Apochromat 25×/0.8 Imm autocorr FCS M27 objective was used. For high-resolution imaging, either a C-Apochromat 40×/1.2 W autocorr FCS M27 water-immersion objective or a Plan-Apochromat 63×/1.4 Oil DIC M27 objective was employed. Depending on the fluorophores, laser lines at 405□nm, 488□nm, 561□nm, or 633□nm were used for excitation.

### Kaede mRNA Synthesis and injection

Kaede mRNA was synthesized using the HiScribe™ T7 ARCA mRNA Kit with poly(A) tailing (New England Biolabs, E2060S) from a PCR-amplified template of the Kaede-H2B plasmid (Addgene, #57316). Following in vitro transcription, the mRNA was purified using SPRISelect magnetic beads (Beckman Coulter, B23319). The final injection mix contained 200□ng/□L Kaede mRNA and fluorescein isothiocyanate (FITC; Thermo Fisher, 46425) as an injection tracer. The mixture was injected into fertilized Nematostella eggs.

### Photoconversion

Larvae (3 DPF) expressing *Kaede* mRNA or *Dendra2::ColIV* were anesthetized in 7% magnesium chloride and mounted in a round glass-bottom dish (MatTek, #P35G-1.5-14-C). Photoconversion was performed using an Evident Rapp FV3000 confocal microscope with a 375□nm laser to induce conversion. After 3 days post-photoconversion, animals were transferred to microscopy slides, screened for photo-converted patches, and imaged. Laser lines at 488□nm and 561□nm were used to detect the unconverted and converted forms of Dendra2/Kaede, respectively.

### shRNA Synthesis and Injection

shRNAs were designed using the siRNA Wizard tool (*6*) (Invivogen), and primers were synthesized by IDT. Primer annealing was carried out at 98□°C for 5 minutes, followed by passive cooling to room temperature. In vitro transcription was performed using the T7 MegaShortScript Kit (Invitrogen, AM1354) with a 6-hour incubation. RNA was purified using SPRISelect magnetic beads (Beckman Coulter, B23319) in 46% isopropanol. Samples were incubated at room temperature for 15 minutes, placed on a magnetic stand for 5 minutes, and washed twice with 80% ethanol. After brief drying, RNA was eluted in RNase-free water, aliquoted, and stored at −80□°C. Fertilized eggs were injected with 500– 1500□ng/□L of each shRNA (see Table 2), together with Texas Red–labeled Dextran (ThermoFisher, D3328) as an injection tracer. Microinjections were performed using a FemtoJet Express system (Eppendorf). Injected embryos were maintained at room temperature and transferred to 27□°C on the following day for further development.

**Table 2:**
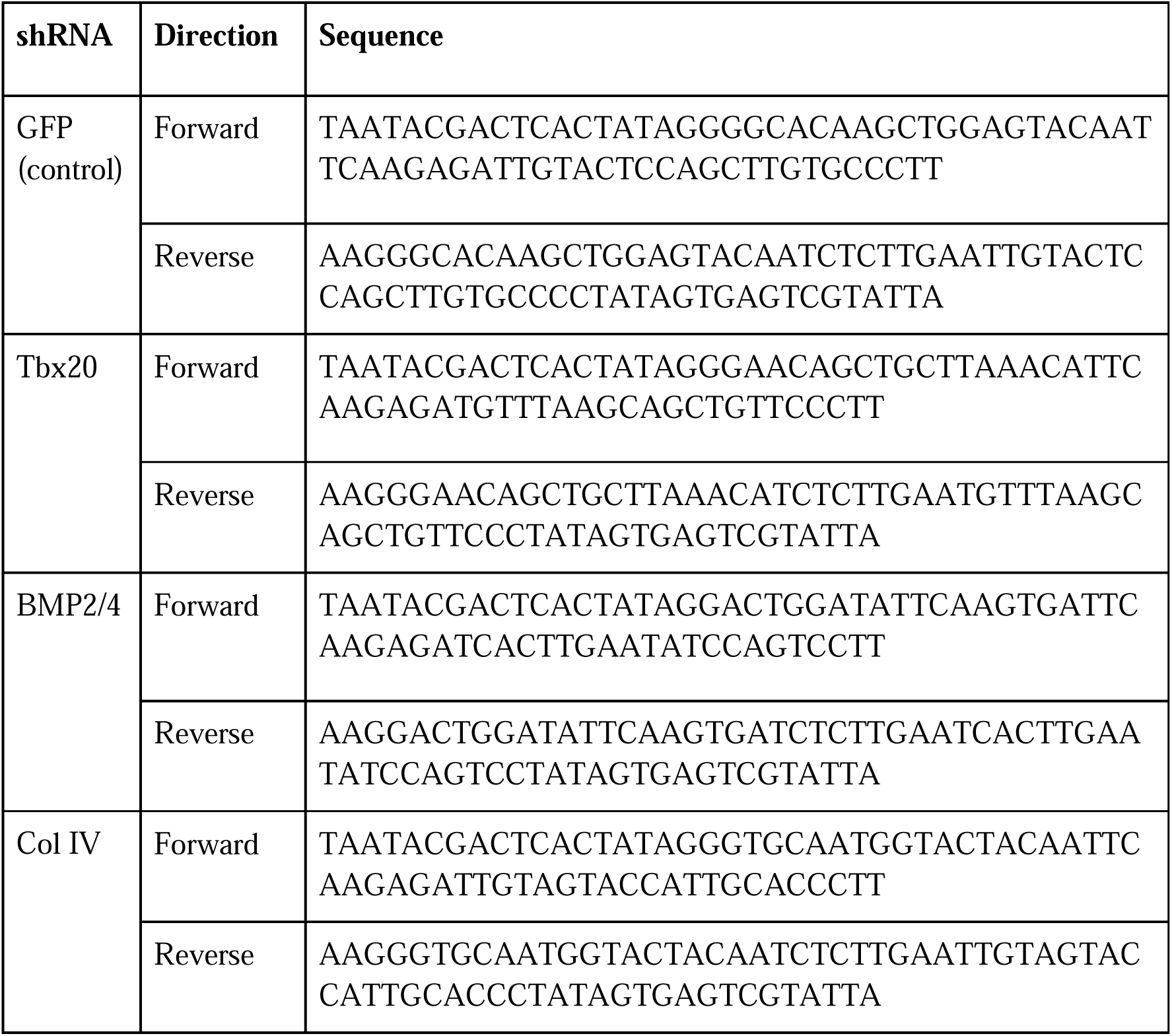
Primer sequences used for shRNA synthesis.

### Pharmacological inhibitor treatments

Larvae at 3 dpf were incubated for 72 hours at 27□°C in 12 ppt ASW containing either 0.1% DMSO (vehicle control), 50□µM GM6001 (Abcam, ab120845), a broad-spectrum matrix metalloproteinase inhibitor, or 50□µM 2,2□-Bipyridine (BPY) (Sigma, 1030980005), an inhibitor of prolyl-4-hydroxylase or 0.5 mM Rocuronium bromide, a previously characterized muscle relaxant (*5*). Drug solutions were prepared fresh and used without replacement over the course of the incubation.

### Quantifications

Image analysis was performed with *Fiji* (*7*) with its *MorphoLibJ* package (*8*). PyBoat in Python was used for time series analysis (*9*). Data analysis and plotting was performed with R in RStudio environment (*10*), with the following packages: tidyverse (*11*), ggplot2 (*12*) and ggsignif (*13*).

#### Morphometric measurements

Confocal cross-sections at the oral plane were used, which is a good approximation given the radial symmetry of the body plan. Only the body column was used for measurements, ignoring the tentacles. To obtain the shape metrics, a bounded box was fitted on the body column, the dimensions of which provided body length *l* (parallel to oral-aboral axis) and *w* (perpendicular to oral-aboral axis). The aspect ratio was used as a metric of shape, which was defined as *l/w*. The aspect ratio correlates strongly with the developmental time, since the morphogenesis is characterized by axial elongation. A polyline ROI was defined along the mesoglea of the body column denoting axial position (0 towards aboral pole, 1 towards oral pole). Using the ROI the region was straightened, which defined a new direction perpendicular to the body axis, where the mesoglea was denoted as 0, positive value (above) denotes the endoderm while negative value (below) denotes the ectoderm. The tissue thickness is reported to be the average thickness along this straightened axis. For eGFP::ColIV+ animals, the eGFP intensity peaked close to 0, the full-width half-maximum (FWHM) value of which denoted the local mesoglea thickness while the signal within denoted the local ColIV amount. The mesoglea thickness and intensity reported were the mean and standard deviation across the entire body column. The same was done for immunostaining (Fig S1B).

#### Aspect ratio of the photo-converted patches

The photo-converted areas *A* were segmented and its skeleton was defined as *l*. The width was defined empirically as *A/l*, and the reported *AR_patch_ = l/w = l^2^/A*. If the shape inverted during axial elongation, the AR was inverted.

#### ECM intensity measurements in perturbations

The intensity of the ECM was measured on maximum intensity projection images by taking the average intensity normalized over equal sized square ROIs.

#### pErk intensity measurements

pErk intensity was measured at the aboral pore, 0µm denoting outside and 50µm denoting the body cavity. The profile plot was generated across a line of width 5µm.

#### Intensity measurements at the aboral end

A ROI of length 100µm was defined along the aboral pore and rescaled, with 0 denoting the location of the pore. The intensity was measured as profile plots.

### Cavity inflation assay

To artificially increase internal pressure, a glass needle containing fluorescein isothiocyanate (FITC; Thermo Fisher, #46425, diluted 1:100 in 12□ppt ASW) was gently inserted into the oral opening of the animal. Injection was carried out using a FemtoJet Express microinjector (Eppendorf), while monitoring the animal under a dissection microscope. Cavity inflation was gradually induced by precisely controlling both the injection pressure and duration. Time-lapse recordings were acquired at 1-second intervals and continued until leakage was observed through either the aboral or oral poles.

### FIB-SEM volume acquisition

The samples were high pressure frozen in a solution of sea water containing 20% Ficoll (molecular weight ∼70,000, Sigma) using HPM010 (Abra Fluid). Following high-pressure freezing, freeze-substitution was carried out using the EM-AFS2 system from Leica Microsystems. The freeze-substitution medium consisted of 0.1% uranyl acetate in acetone. The samples were subjected to freeze-substitution at a temperature of -90°C for a duration of 48 hours. Subsequently, the temperature was gradually increased to -45°C at a rate of 3.5°C per hour, and the samples were further incubated for 5 hours. The samples were gradually infiltrated in HM20 resin, then polymerized under UV light for a period of 48 hours at a temperature of -25°C. Following this, the temperature was gradually increased to 20°C at a rate of 5°C per hour, and the samples were further UV polymerized for an additional 9 hours. In order to target the central aboral region of the animal with sufficient accuracy for FIB-SEM acquisition, we used an already established strategy (*14*). The samples, mounted on SPINE sample holders commonly used for crystallography, were imaged by phase-contrast X-ray on the EMBL beamline P14 on the PETRA III synchrotron (c/o DESY, Hamburg, Germany) using a previously characterized imaging setup (*15*) at an X-ray energy of 18 keV. X-ray images were recorded using an Optique Peter (Lyon, France) X-ray microscope consisting of an LSO:Tb scintillator with 8 □m active layer; an Olympus UPlanFL 20-fold objective (Olympus, Tokio, Japan), numerical aperture 0.5; a 45° mirror; a 180 mm tube lens and a PCO.edge 4.2 sCMOS camera with 2048x2048 pixels, pixels (6.5 □m pixel size). Thus, the effective pixel size was 0.325 □m with a field of view 666 x 666 □m². This setup typically delivers a resolution of about 0.5-0.7 □m, as determined from the analysis of projection images from a Siemens star (Ta on SiN; XRESO-50HC, NTT-AT, Japan). On the resin-embedded sample, projection images were acquired at four camera distances: 62.5, 67.5, 73.5 and 82.5 mm. At each distance, 3600 projections covering 360° of continuous rotation were recorded with an exposure time of 10 ms per frame. Data collection (including robotic sample transfer from a storage vessel to the rotation axis and sample centering via an on-axis optical microscope) was completed in 6 minutes. Flat-field corrections were applied by dividing each projection image by the most similar flat-field image according to the SSIM criterion (*16*). For lateral shift compensation at the four camera positions, images recorded at each projection angle were registered using Fourier-space correlation with a sub-pixel interpolation. Registered images were further processed by a multi-distance non-iterative holographic reconstruction (*17*, *18*), using a complex refraction index decrement ratio β/δ = 0.15 and a zero compensation of 0.1. Tomographic reconstructions were performed using the TOMOPY package (*19*), employing the built-in Gridrec algorithm and Shepp-Logan filtering with default settings. All steps of the XIMG data processing were combined into a python-based custom software pipeline, available at:

https://git.embl.de/maxim.polikarpov/ximg_p14/-/blob/master/2019/Dec_2019_Platynereis/nematosella/

The high contrast, high resolution X-ray data, were used to guide the trimming of the block to the region of interest, which was done with an ultramicrotome (UC7, Leica Microsystems) and a diamond trimming knife (Cryotrim 90, Diatome). The trimmed sample was then glued to an SEM stub using conductive epoxy resin (Ted Pella) and imaged by FIB-SEM using a Zeiss Crossbeam 550. The acquisition was performed using the Atlas 3D workflow. FIB slicing was obtained at 1.5 nA. Imaging was performed at an acceleration voltage of 1.5 kV and a current of 700pA, using a backscattered electron detector (ESB). The voxel size was 16x16x16 nm, with a dwell time of 10 µs.

### FIB-SEM volume segmentation

Cell segmentation from FIB-SEM data was performed using a boundary-based semantic segmentation pipeline adapted from (*20*). Initial boundary annotations were generated in *Ilastik* (*21*), which was used to train a random forest classifier for preliminary segmentation.

This segmentation was subsequently refined using a publicly available pretrained model (https://bioimage.io/#/?tags=enhancer&id=10.5281%2Fzenodo.6808325) applied as an enhancer. To further improve accuracy, a U-Net architecture (*22*) was trained from scratch using the enhanced intermediate segmentation as training labels. Following the protocol in (*20*), the final boundary map was converted into an instance-level segmentation using a watershed algorithm (*23*). Minor segmentation artifacts were manually corrected before rendering the target cells for visualization.

