## Supplemental Figures for "Mesoglea biogenesis reveals a cryptic aboral valve for pressure regulation in cnidarian morphogenesis"

**Figure S1 to S7**

**Movies S1 to S7**

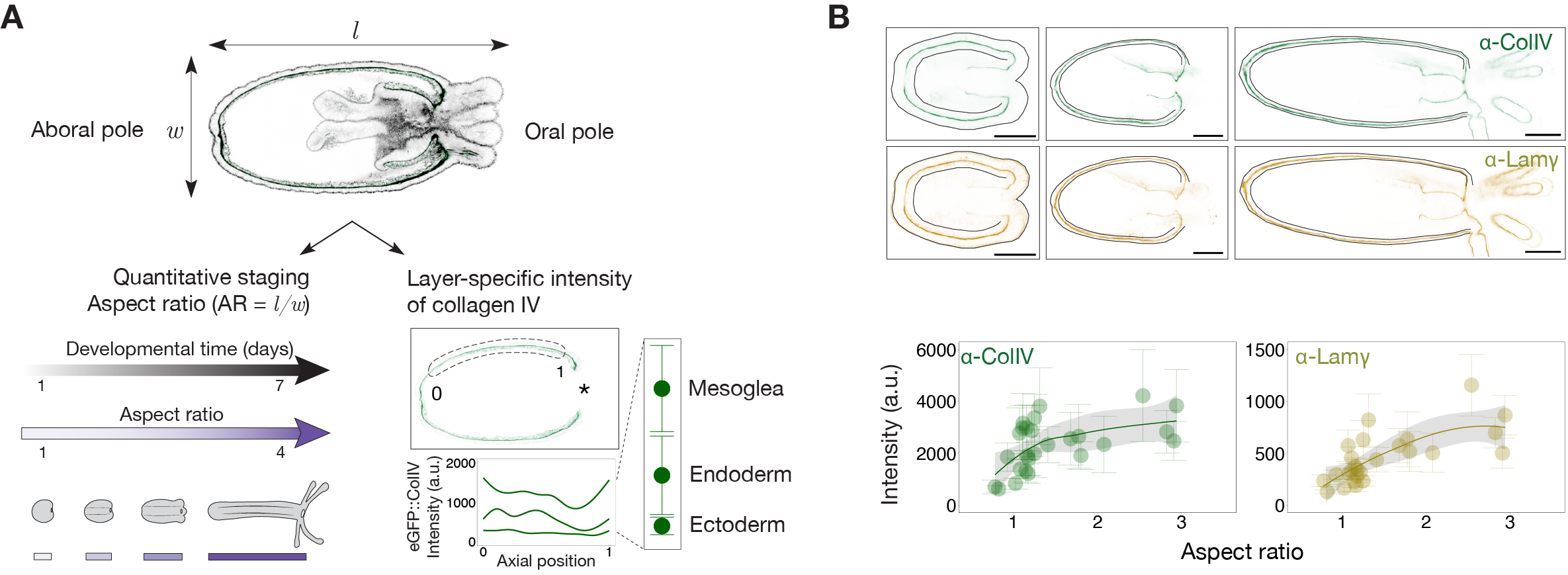

**Figure S1: Quantification of Col IV and Laminin**

(**A**) Schematic of developmental morphometrics and the regions used for quantifying *eGFP::ColIV* intensity.

(**B**) (Top) Confocal cross-sections of animals immunostained for Collagen IV and Laminin across developmental stages. Scale bar: 100µm. (Bottom) Quantification of extracellular Collagen IV and Laminin intensities as a function of body aspect ratio.

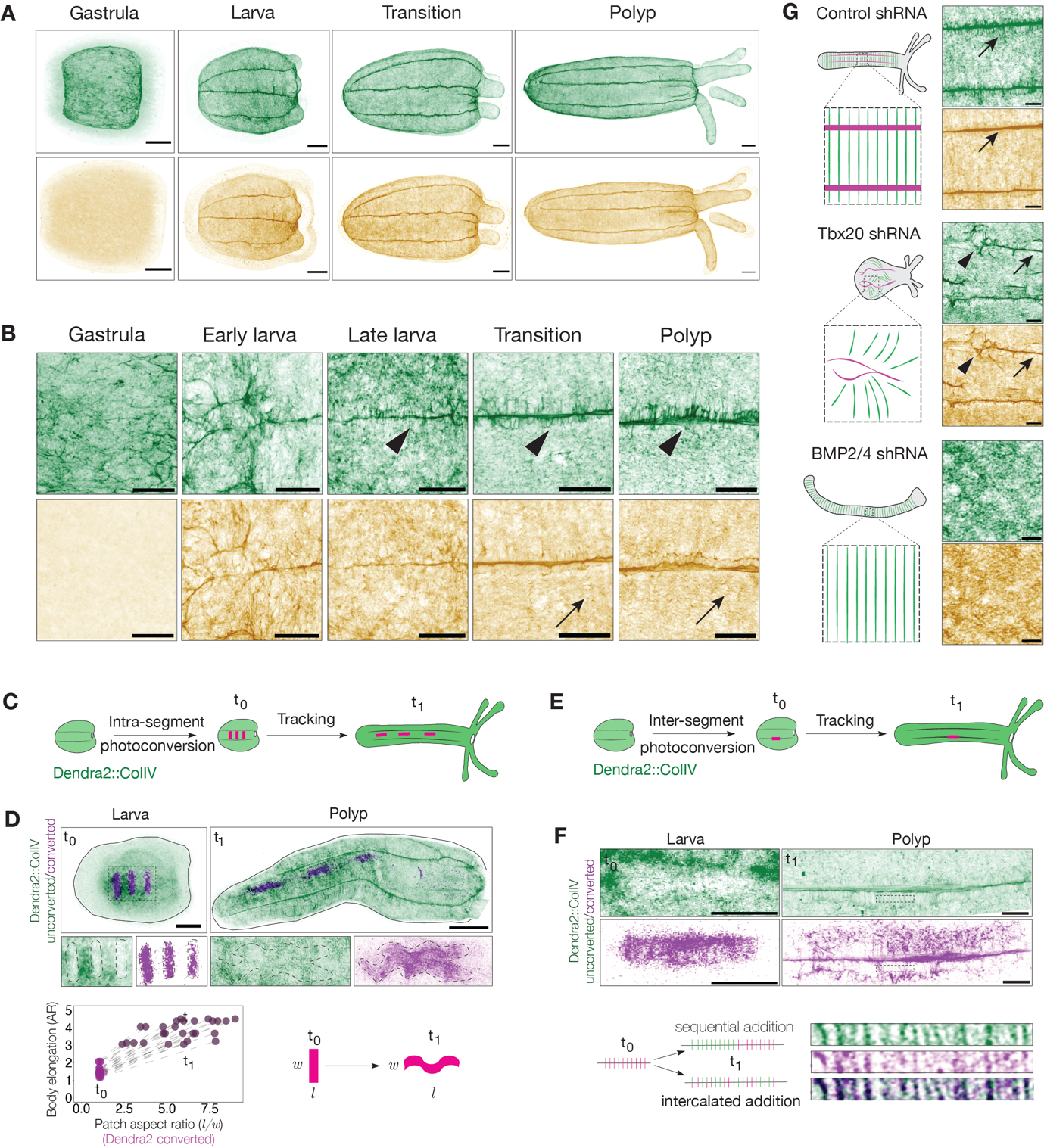

**Figure S2: developmental dynamics of Collagen IV and Laminin**

(**A**) Maximum intensity projections showing Collagen IV and Laminin immunostaining across developmental stages. Scale bar: 50µm.

(**B**) Magnified views of the regions shown in A, highlighting the spatial organization of Collagen IV and Laminin. Scale bar: 20µm.

(**C**) Schematic of the Dendra2::ColIV photoconversion assay in intra-segmental regions. Green: non-photoconverted; magenta: photoconverted.

(**D**) Time-course of photoconverted intra-segmental patches (magenta) from larva to primary polyp (*n =* 7 animals). Upper: Photoconverted regions at t_0_ (day of conversion) and t_1_ (3 days post-conversion). Lower: Zoom-in views of corresponding patches. Plot shows aspect ratio changes of tracked patches as a function of overall body elongation. Schematic summarizing the fate of intra-segmental Dendra2::ColIV photoconverted regions across development. Scale bar: 50 µm.

(**E**) Schematic of the Dendra2::ColIV photoconversion experiment in inter-segmental regions.

(**F**) Photoconverted inter-segmental patches (magenta) tracked from larva to primary polyp (*n =* 5 animals). Zoom-in shows the emergence of lateral Collagen IV bridges. Two models for lateral fiber addition are shown. Scale bar: 20 µm.

(**G**) Gene knockdown experiments disrupt endoderm morphogenesis and muscle organization (green: circular muscles; magenta: longitudinal muscles), showing resulting patterns of Collagen IV (green) and Laminin (yellow). *n =* 6 animals for control shRNA; *n =* 6 animals for Tbx20 shRNA; *n =* 6 animals for BMP2/4 shRNA. Scale bar: 20 µm.

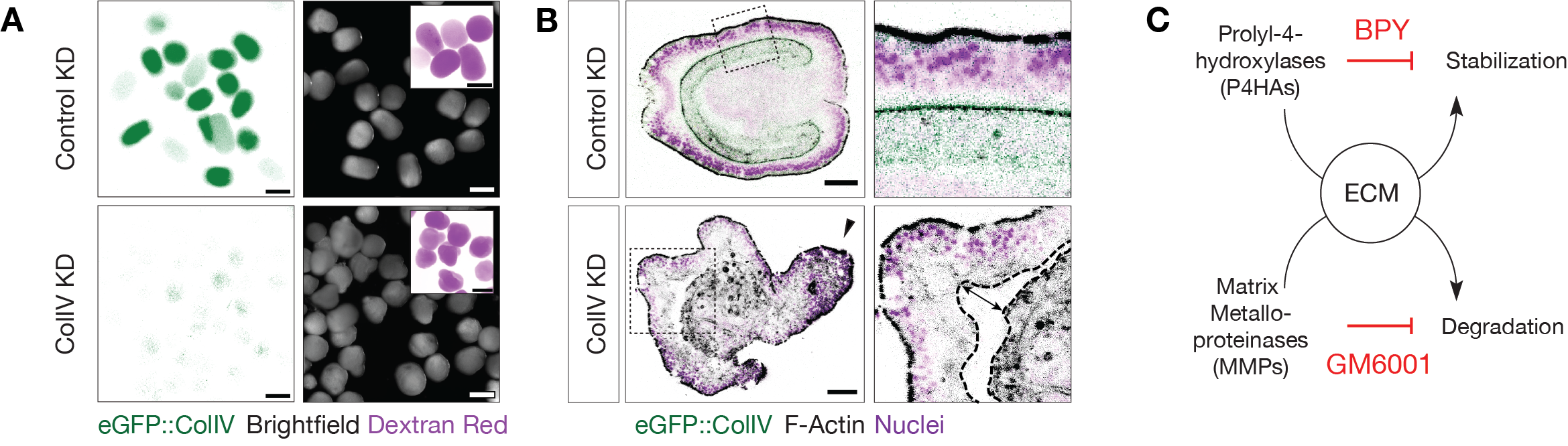

**Figure S3: Loss of function of Collagen IV and pharmacological modulation of ECM**

**(A)** Epifluorescence and brightfield images of larvae at 3 DPF from an *eGFP::ColIV* heterozygous cross with wild type, following control or *ColIV* shRNA injection. In control knockdown (KD) animals show normal morphology (*n =* 20), with ~50% of larvae expressing *eGFP::ColIV* as expected. In contrast, *ColIV* KD animals display a marked reduction in *eGFP::ColIV* signal and abnormal morphology (*n =* 47/51). Scale bar: 200 µm.

**(B)** Confocal cross-sections of representative control and *ColIV* KD larvae stained for F-actin and Hoechst (*n =* 8 each). Note the disorganized tissue architecture (arrowhead) and pronounced gap between ectoderm and endoderm (double-headed arrow) in the Col IV KD condition. Scale bar: 50 µm.

**(C)** Schematic summarizing pharmacological inhibitors targeting ECM stabilization and degradation.

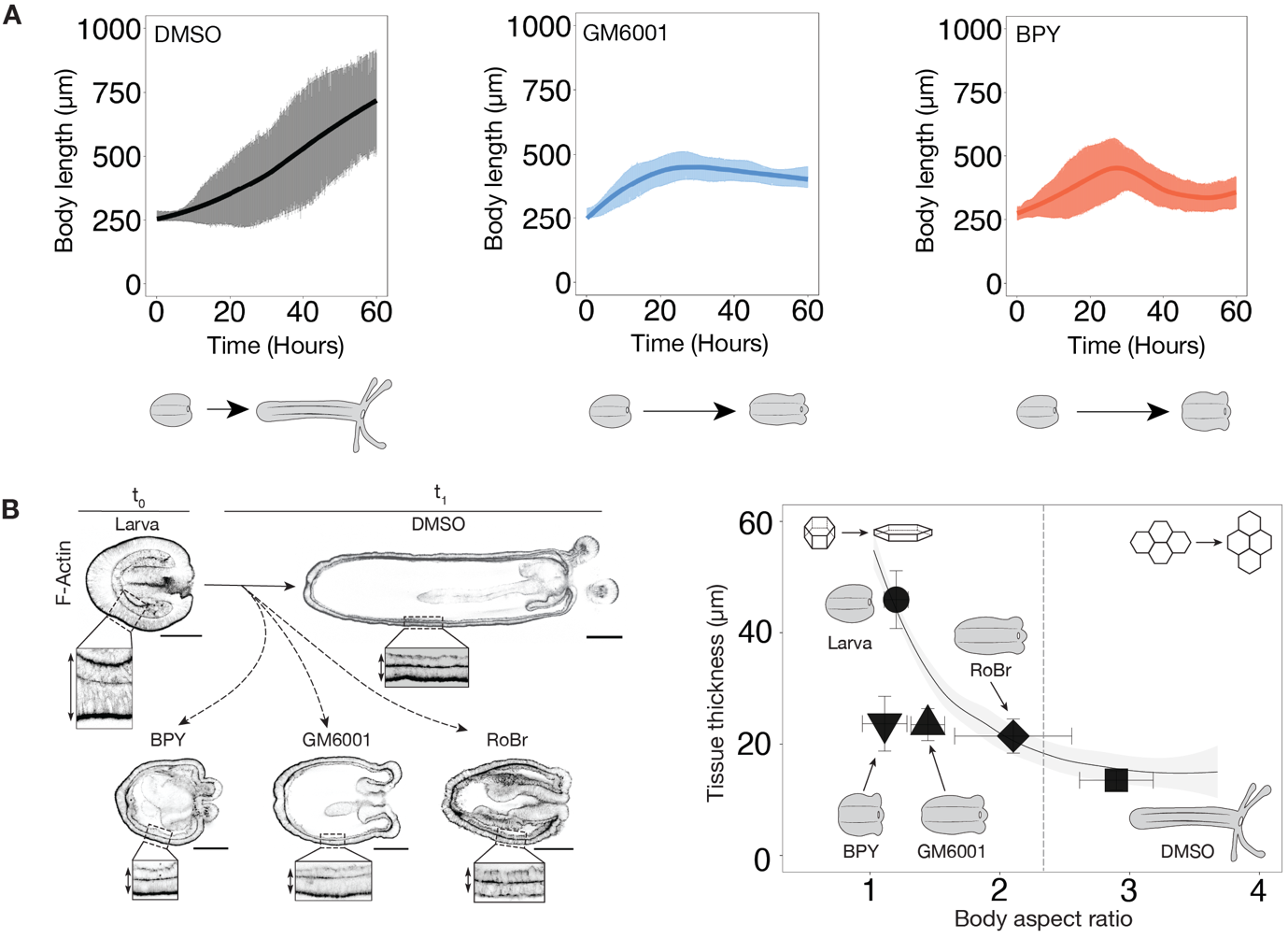

**Figure S4: Elongation dynamics in control and drug-treated animals**

(**A**) Plots showing changes in body length over time for each treatment condition. Solid lines represent the mean, and shaded areas indicate the standard deviation.

**(B)** (left) Mid-plane optical sections of animals stained for F-actin. Zoom-in views highlight alterations in body wall architecture under each treatment condition. Scale bar: 100µm. (Right) Quantification of body wall thickness as a function of body aspect ratio across all conditions. The black curve and light gray shading represent the mean and standard deviation of thickness measurements during normal development. *n* = 20 DMSO, *n* = 20 BPY, *n* = 20 GM6001, and the muscle anesthetic Rocuronium bromide (*n* = 20 RoBr).

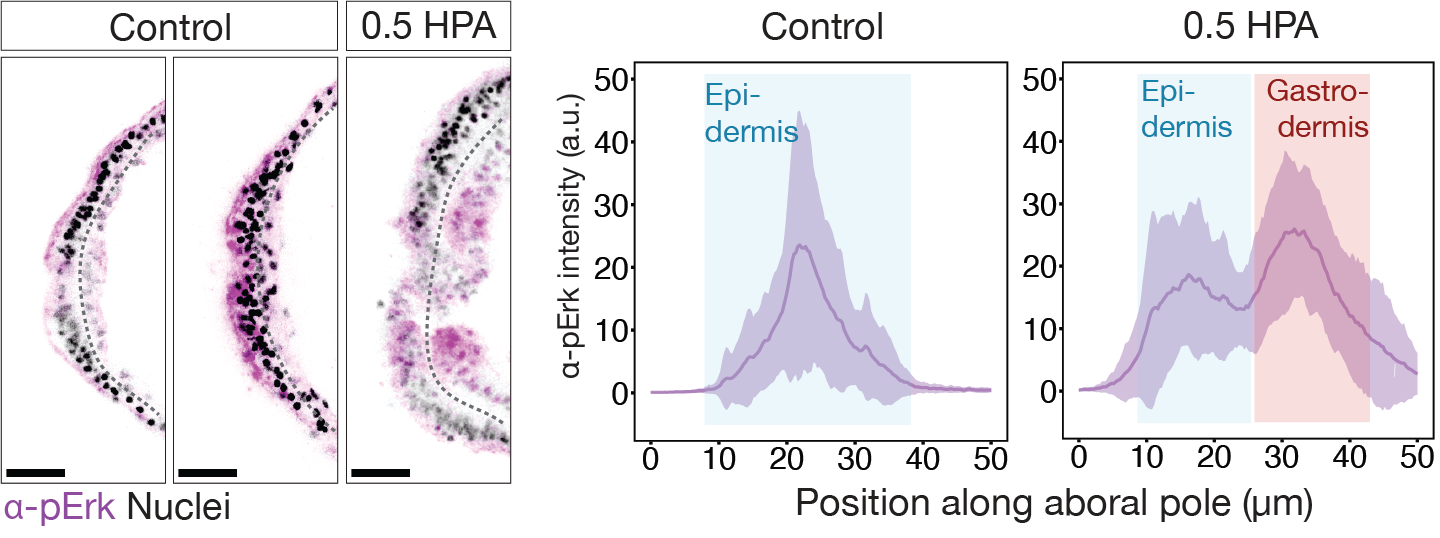

**Figure S5**: **Activity of ERK signaling at the aboral pole.**

(Left) Cross-sectional images of control polyps stained for pERK and Hoechst, showing two representative patterns of pERK signal intensity. Additional panel shows pERK staining 0.5 hours post-ablation (HPA) of the aboral pole. Scale bar: 20 µm. (Right) Quantification of pERK intensity in control versus ablated animals. Purple curve and light shading represent the mean and standard deviation. Signal localization within the epidermis versus endodermis is indicated.

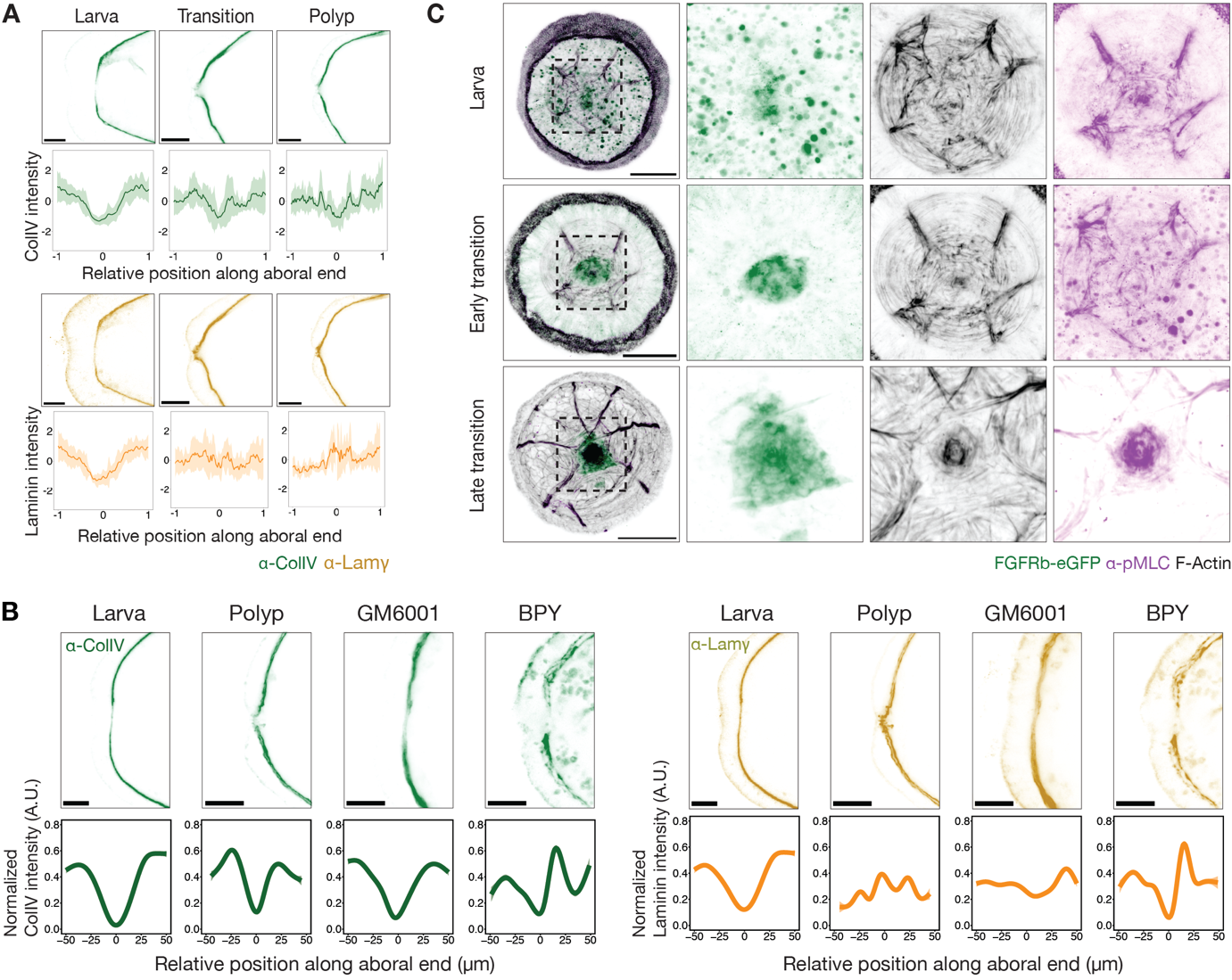

**Figure S6: Morphogensis of the aboral valve and effect of pharmacological inhibitors on ECM organisation at the aboral pole.**

**(A)** Cross-sectional images showing immunostaining for Collagen IV and Laminin at the aboral end across the indicated developmental stages. Plots show the spatial distribution of Collagen IV and Laminin intensities at each stage, *n =* 6 for each; solid lines represent the mean, and shaded areas indicate the standard deviation. Scale bar: 10 µm.

(**B**) Cross-sectional images showing Collagen IV and Laminin immunostaining at the aboral end across the indicated treatment conditions. Plots show the spatial distribution of Collagen IV and Laminin intensities at each condition; solid lines represent the mean. Scale bar: 10 µm.

**(C)** Aboral views of *FGFRb-eGFP* animals stained for F-actin and pMLC across the indicated developmental stages. Scale bar: 20µm.

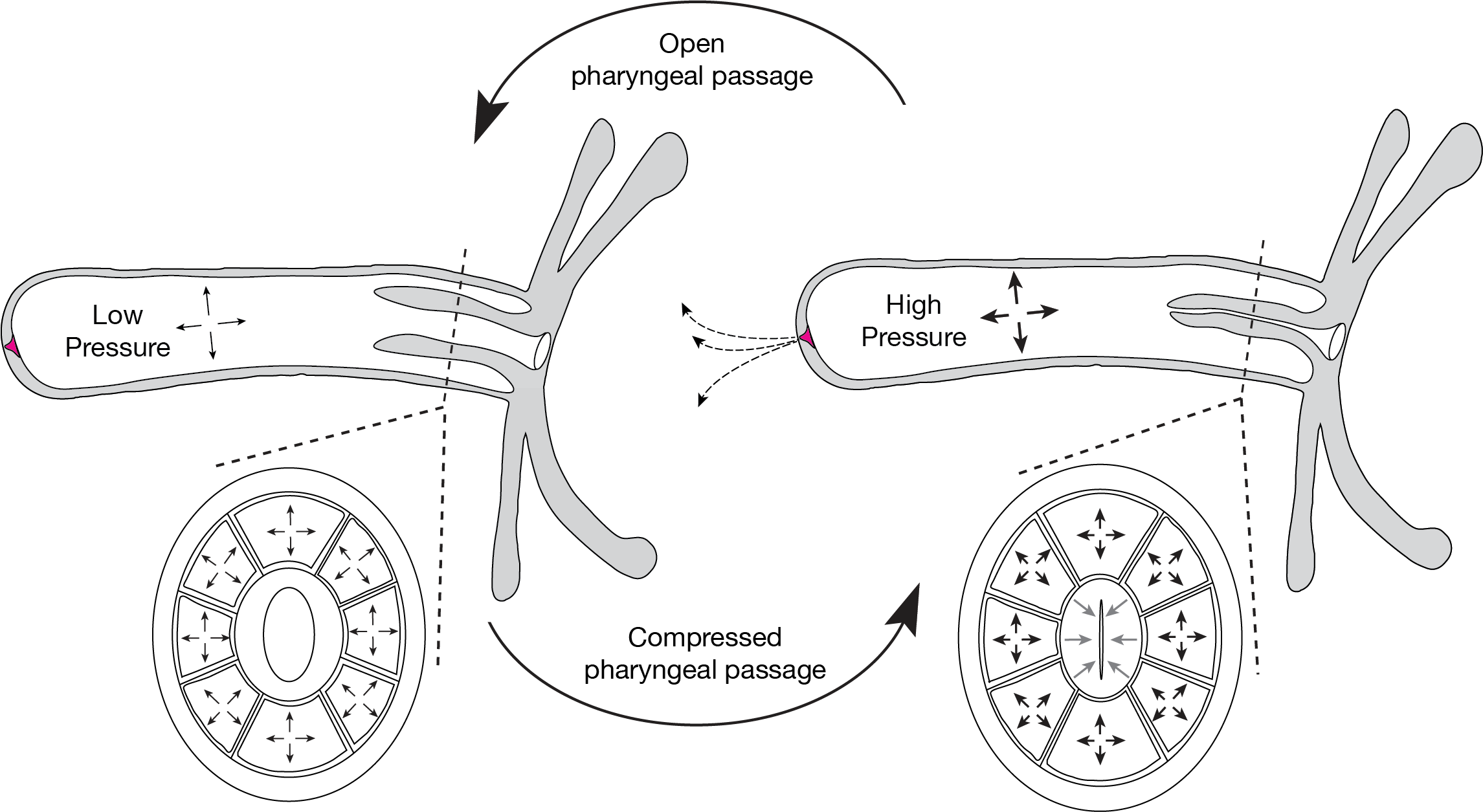

**Figure S7. Working model of pharyngeal states under low and high internal cavity pressure.** The schematic illustrates a primary polyp and a cross-section of the pharyngeal region. Note that the pharynx is connected to the body wall via eight mesenteries, forming eight fluid-filled pouches that are directly continuous with the main body cavity. Under low internal pressure, the pharyngeal passage remains open, permitting potential flow through the oral opening. In contrast, elevated pressure within these pouches compresses the pharynx, effectively sealing the passage and preventing fluid release through the oral end. Under this high-pressure scenario, the aboral opening functions as a release valve, enabling pressure relief through the aboral end.

**Movie S1:** Live imaging of early eGFP::Col IV embryos using confocal and light-sheet microscopy. Part 1: confocal time-lapse of multiple embryos undergoing gastrulation. Part 2: lightsheet time-lapse of a single gastrulating embryo.

**Movie S2:** Live imaging of larva-polyp transition under the indicated conditions. Part 1: DMSO-control, Part 2: GM6001-treated and Part 3: BPY-treated animals.

**Movie S3**: Live imaging of aboral leakage. Part 1: DMSO-control and Part 2: BPY-treated animals.

**Movie S4.** Segmented FIB-SEM data of the aboral pole**.** Part 1: XY view, with oral side at the top and aboral side at the bottom. Note the unique structure of the mesoglea visible at 00:08. Part 2: XZ view, progressing from the epidermis to the gastrodermis. Observe the basal epidermal gap beginning at 00:17.

**Movie S5**: 3D rendering showing the dual conformational states of the muscular valve labeled with *FGFRb-eGFP*. Part 1: closed state, Part 2: open state and Part 3: Localisation of pMLC at the muscular valve.

**Movie S6**: Live imaging of cavity inflation during development. Part 1: larva, Part 2: early transition, Part 3: late transition and Part 4: primary polyp.

**Movie S7**: Live imaging of cavity inflation in the FGFRb mutant and its sibling. Part 1: a sibling of the FGFRb mutant (control), Part 2: early response of the FGFRb mutant, and Part 3: late response of the FGFRb mutant.
